# Breaking the Extraction Bottleneck: A Single AI Agent Achieves Statistical Equivalence with Human-Extracted Meta-Analysis Data Across Five Agricultural Datasets

**DOI:** 10.64898/2026.02.17.706322

**Authors:** Moshe Halpern

## Abstract

**Background:** Data extraction is the primary bottleneck in meta-analysis, consuming weeks of researcher time with single-extractor error rates of 17.7%. Existing LLM-based systems achieve only 26–36% accuracy on continuous outcomes, and no study has validated AI-extracted continuous data against multiple independent datasets using formal equivalence testing.

**Methods:** A single AI agent (Claude Opus 4.6) extracted treatment means, control means, sample sizes, and variance measures from source PDFs across five published agricultural meta-analyses spanning zinc biofortification, biostimulant efficacy, biochar amendments, predator biocontrol, and elevated CO2 effects on plant mineral nutrition. Observations were matched to reference standards using an LLM-driven alignment method. Validation employed proportional TOST equivalence testing, ICC(3,1), Bland-Altman analysis, and source-type stratification.

**Results:** Across five datasets, the agent produced 1,149 matched observations from 136 papers. Pearson correlations ranged from 0.984 to 0.999. Proportional TOST confirmed statistical equivalence for all five datasets (all p < 0.05). Table-sourced observations achieved 5.5x lower median error than figure-sourced observations. Aggregate effects were reproduced within 0.01–1.61 pp of published values. Independent duplicate runs confirmed extraction stability (within 0.09–0.23 pp).

**Conclusions:** A single AI agent achieves statistical equivalence with human-extracted meta-analysis data across five independent agricultural datasets. The approach reduces extraction cost by approximately one to two orders of magnitude while maintaining accuracy sufficient for aggregate meta-analytic pooling.

**Highlights:** *What is already known:*

- Data extraction is the primary bottleneck in meta-analysis, with single-extractor error rates of 17.7%
- Existing LLM-based extraction systems achieve only 26-36% accuracy on continuous outcomes
- No study has validated AI extraction against multiple independent datasets using formal equivalence testing

*What is new:*

- A single AI agent achieves statistical equivalence with human-extracted data across five agricultural meta-analyses (1,149 observations, 136 papers)
- LLM-driven alignment resolves the previously underappreciated bottleneck of moderator matching, improving correlations from 0.377-0.812 to 0.984-0.997 without changing extracted values
- Table-sourced observations achieve 5.5x lower error than figure-sourced data

*Potential impact for RSM readers:*

- Provides a validated, reproducible workflow for AI-assisted data extraction in meta-analysis
- Demonstrates that most apparent “extraction error” in validation studies is actually alignment error
- Offers practical quality signals (source-type labeling) for downstream meta-analysts

## 1. Introduction

### 1.1 The Extraction Bottleneck

Meta-analysis is central to evidence-based practice in agricultural science, yet the systematic review process remains both slow and fragile. The mean time to complete and publish a systematic review is 67.3 weeks ^1^, and 23% of reviews need updating within two years of publication ^2^. The primary bottleneck is not statistical analysis but data extraction: trained researchers must manually identify, read, and record quantitative values from source publications.

This bottleneck is expensive and error-prone. Manual extraction requires an estimated 2–8 hours per paper ^3^. Single-extractor error rates reach 17.7%, falling to 8.8% only under costly dual-extraction protocols ^4^ – underpinning the Cochrane Handbook’s recommendation for dual independent extraction ^5^. At the meta-analysis level, 66.8% of published meta-analyses contain at least one data extraction error ^6^. For agricultural meta-analyses spanning hundreds of papers with complex factorial designs, the extraction burden can extend to months of researcher time.

### 1.2 Challenges in Agricultural Data

Agricultural experiments present distinct challenges for automated extraction. Lacking standardized reporting frameworks like CONSORT ^7^, plant science studies employ complex factorial designs (e.g., CO2 x cultivar x soil amendment x harvest date), producing multi-layered tables where only specific treatment combinations are relevant. Variance reporting is inconsistent: nearly 70% of ecological meta-analysis datasets include studies with missing standard deviations ^8^. Effect sizes span orders of magnitude – from single-digit percentage changes in mineral concentrations to >200% increases in zinc biofortification studies – complicating any uniform accuracy threshold.

### 1.3 The State of LLM-Based Extraction

Recent progress in large language models (LLMs) has opened new possibilities for automating data extraction, but results have been mixed. For categorical variables (study design, population characteristics), LLMs consistently achieve high accuracy: Gougherty and Clipp^9^ reported >90% accuracy for ecological categorical data, and Helms Andersen et al^10^ found F1 scores around 90% for Cochrane data extraction. However, performance drops substantially for continuous numerical outcomes. Jansen et al^11^ evaluated GPT-4o for extracting means, standard deviations, and sample sizes from psychology reviews, reporting 26–36% accuracy. Kataoka et al^12^ tested o3 for clinical data extraction, achieving 75.3% overall accuracy but concluding that numeric extraction was “still inadequate” for unsupervised use.

Several systems have demonstrated stronger performance under specific conditions. Cao et al^13^ developed OttoSR, achieving 93.1% accuracy on structured clinical data extraction across 7 Cochrane reviews. Gartlehner et al^14,15^ found AI-assisted extraction achieved 91.0% accuracy versus 89.0% for human-only extraction, with a median time saving of 41 minutes per study. Poser et al^16^ showed that three-model consensus reduced extraction errors to 1.48% for clinical reports. Khan et al^17^ demonstrated that LLMs in a collaborative two-reviewer workflow outperformed individual LLMs. However, virtually all published validation studies operate on clinical or medical data – randomized controlled trials, Cochrane reviews, and CONSORT-formatted reports. The sole exception outside medicine is Gougherty and Clipp^9^ in ecology, which found LLMs poor at quantitative extraction (23.8% accuracy using GPT-3.5).^18,19^ The agricultural domain, with its complex factorial designs and non-standardized reporting, remains largely untested.

### 1.4 Three Underappreciated Gaps

Beyond the overall accuracy question, three critical gaps persist in the literature. First, no study has applied formal equivalence testing (e.g., TOST) to AI-extracted continuous data. Demonstrating low error is necessary but not sufficient; formal statistical equivalence is required to justify substituting AI for human extraction. Second, no study has validated across multiple independent ground-truth datasets. Single-dataset validation cannot distinguish genuine capability from dataset-specific overfitting. Third, no study has separated extraction error from alignment error – the distinction between reading wrong values from a PDF versus matching correct values to the wrong reference-standard row.

A further challenge is epistemic circularity: if the same ground truth used during system development also serves as the validation benchmark, performance estimates may be inflated. Breaking this circularity requires true holdout datasets and independent replication runs that demonstrate extraction stability.

### 1.5 Study Aims

We present a validation study of a single AI agent extracting quantitative data from scientific PDFs across five published agricultural meta-analyses. We address four questions:

1. Does AI-extracted data achieve formal statistical equivalence with published reference standards?
2. Does this equivalence hold across diverse agricultural domains and effect-size scales?
3. How much apparent “extraction error” is actually alignment error?
4. Is extraction stable across independent runs?

To our knowledge, this is the first study to (a) formally test equivalence of AI-extracted continuous data against published reference standards using proportional TOST, (b) validate across five independent datasets spanning diverse agricultural domains, (c) demonstrate extraction stability through independent duplicate runs, and (d) quantify the contribution of alignment error to apparent extraction error.

## 2. Methods

### 2.1 Agent Architecture

A single AI agent (Claude Opus 4.6, Anthropic, running within the Claude Code CLI environment) read each source PDF directly and extracted observations. We use the term “agent” to denote an AI system that autonomously reads documents and produces structured output – specifically, a system that receives a PDF and a natural-language instruction, then outputs a structured JSON file with one object per observation.

The agent operated within a 200K-token context window, sufficient to ingest most scientific papers in their entirety. For each paper, the agent received a brief natural-language instruction specifying the treatment, control, outcome variable, and desired output fields (full prompts in Appendix A). The agent produced structured JSON following a predefined schema specifying fields for paper metadata, experimental conditions, treatment and control means, sample size, variance measures, and moderator variables. No domain-specific prompt templates, few-shot examples, or vision extraction pipelines were used. The same agent model was used for all five datasets; only the natural-language instruction varied.

#### 2.1.1 Three-Interface Architecture

The extraction system is deployed through three interfaces to accommodate different use cases (Table 1).

**TABLE 1:** Three deployment modes of the extraction system.

| Mode | Interface | Requirements | Use case |
| --- | --- | --- | --- |
| **Skill** | Claude Code slash command | Claude Code CLI | Interactive single-paper extraction |
| **MCP Server** | 7 tools via Model Context Protocol | Claude Desktop + API key | GUI-based extraction workflow |
| **Standalone CLI** | Command-line script | API key only | Batch processing, automation |

All modes share the same core extraction pipeline. The MCP (Model Context Protocol) server exposes extraction, validation, and export as individual tools callable from Claude Desktop or any MCP-compatible client. This architecture allows researchers to interact with the system through whichever interface matches their technical comfort and workflow requirements.

#### 2.1.2 Source Type Labeling

Every extracted observation was labeled with its data source type: **table** (numeric data read from structured tables), **figure** (values estimated from bar charts, scatter plots, or other graphical displays), or **text** (numbers cited in running prose). This labeling provides transparent quality flags, since extraction accuracy differs substantially by source type (Section 3.4).

#### 2.1.3 Extraction Workflow

For all five datasets, a three-pass extraction workflow was employed: (1) initial extraction from the full PDF, (2) text cross-check comparing extracted values against original PDF text to flag potential errors, and (3) targeted re-extraction of flagged observations. For the Loladze 2014 dataset, additional extraction passes were used to maximize observation coverage across the 23-element, multi-factorial design.

#### 2.1.4 Computational Requirements

Single-paper extraction required approximately 2–5 minutes of processing time. Using the Claude Opus 4.6 API ^20^ ($5 per million input tokens, $25 per million output tokens), each paper consumed approximately 80,000 input tokens and 8,000 output tokens, yielding a per-paper cost of approximately $0.60. The complete extraction of all five datasets (∼179 papers), including alignment and variance recovery passes, cost approximately $150–250 in total API fees and required approximately 6–8 hours of unattended processing time. These estimates exclude the time for prompt development and validation, which was a one-time fixed cost amortized across all datasets.

### 2.2 Validation Datasets

Five published meta-analyses served as reference standards for validation (Table 2). No agent parameters were tuned against any reference standard. All five datasets were processed independently, with the agent receiving only the dataset-specific instruction (treatment definition, control definition, outcome variable).

**TABLE 2:** Validation dataset characteristics.

| Dataset | Domain | Papers processed | Papers matched | Obs matched | Mean effect (%) | Effect range |
| --- | --- | --- | --- | --- | --- | --- |
| Hui 2025 | Zn biofortification / wheat | 37 | 17 | 319 | ~50 | 0–250+ |
| Li 2022 | Biostimulant / crop yield | 50 | 31 | 117 | ~17 | -20 to 80 |
| Li 2024 (biochar) | Biochar / crop yield | 28 | 26 | 254 | ~12 | -30 to 100+ |
| Boldorini 2024 | Predator biocontrol / yield | 18 | 17 | 46 | ~37 | -10 to 100 |
| Loladze 2014 | CO<sub>2</sub> / mineral nutrition | 46 | 45 | 413 | ~-8 | -50 to 50 |
| **Total** |  | **179** | **136** | **1,149** |  |  |

#### Hui et al. 2025 ^21^

A global meta-analysis of zinc agronomic biofortification in wheat and its drivers, published in *Nature Communications*. This dataset features standardized single-element tabular data with consistent reporting formats. Effect sizes are large (mean ∼50%), reflecting substantial zinc concentration increases from biofortification treatments.

#### Li et al. 2022 ^22^

A meta-analysis of non-microbial biostimulant effects on agronomic outcomes, published in *Frontiers in Plant Science*. The raw extraction produced r = 0.453 against reference data, driven entirely by unit-scale mismatches (e.g., yield in t/ha vs. kg/ha). After programmatic harmonization accounting for unit conversions – with no change to any extracted value – the correlation rose to r = 0.994.

#### Li et al. 2024 (biochar yield) ^23^

A meta-analysis of biochar amendment effects on crop yield (2,438 observations, 367 studies), published with open data on figshare. This dataset was processed after all extraction and alignment methods were finalized, serving as a fully prospective holdout. Effect sizes are moderate (mean ∼12%), and 52% of extracted observations came from figures rather than tables.

#### Boldorini et al. 2024 ^24^

A meta-analysis of predator biocontrol effects on crop yield, published in *Proceedings of the Royal Society B*. This dataset extends validation beyond plant nutrition into ecological pest management, with moderate-to-large effect sizes (mean ∼37%) and a small but informative observation set (N = 46).

#### Loladze 2014 ^25^

A comprehensive meta-analysis of elevated CO2 effects on plant mineral concentrations, published in *eLife*. This is the most structurally demanding dataset: complex factorial designs (CO2 x cultivar x ozone x nitrogen), diverse exposure systems (FACE, OTC, growth chambers), and multiple tissues and developmental stages. Effect sizes are small (mean ∼-8%), requiring high extraction precision to detect.

Together, the five datasets span a range of extraction difficulty (simple tabular to complex factorial), effect-size magnitude (8–50 pp), reporting conventions, and scientific domains, providing a diverse testbed for evaluating extraction performance.

### 2.3 Matching Protocol

#### 2.3.1 LLM-Driven Alignment

A key methodological contribution of this work is the use of LLM-driven alignment to match extracted observations to reference-standard rows. Previous approaches relied on either (a) value-based matching, which is circular because it consults the outcome variable, or (b) hardcoded synonym dictionaries (e.g., “corn” = “maize”), which are brittle and incomplete.

Our approach uses an LLM to read both the extracted data schema and the reference-standard schema, then propose moderator mappings automatically. The LLM identifies:

- **Study mappings:** Which extracted paper corresponds to which reference-standard entry (accounting for variant citation styles).
- **Value synonyms:** Equivalent moderator values across naming conventions (e.g., “corn” to “Maize”, “hardwood” to “Wood”).
- **Column mappings:** Which extracted fields correspond to which reference-standard columns.
- **Effect-size format detection:** Whether the reference standard reports raw means, log response ratios, or percentage changes.

The alignment is cached as a human-editable JSON file, enabling manual inspection and correction. On the Li 2024 (biochar) dataset, LLM-driven alignment improved the validation correlation from r = 0.377 (dictionary-based matching) to r = 0.997 with no changes to any extracted value. This improvement demonstrates that apparent “extraction error” can be dominated by alignment failure.

#### 2.3.2 Effect-First Matching for Scale-Heterogeneous Datasets

For datasets where primary studies report outcomes in different units (e.g., crop yield in t/ha, kg/ha, or g/pot), value-based matching on raw means produces spurious mismatches. We developed an effect-first matching approach: observations are first converted to percentage change from control, then matched on derived effect sizes rather than raw values. This approach, combined with automated scale-factor detection (testing common conversion factors: 1000, 100, 10, 0.001, etc.), resolved the Li 2022 correlation from r = 0.453 (raw matching) to r = 0.994 (scale-harmonized matching) without altering any extracted value.

#### 2.3.3 Loladze 2014 Matching

For the Loladze 2014 dataset, the primary validation used LLM-driven alignment (Section 2.3.1), consistent with all other datasets. For comparison purposes (Table 7), we also report results from an earlier metadata-based matching protocol that computed similarity scores across up to 14 dimensions (tissue group, species, site, CO2 level, cultivar, ozone filter, harvest stage, and others) with globally optimal 1-to-1 assignment via the Hungarian algorithm. The substantial improvement from metadata-based to LLM-driven matching (r = 0.812 to 0.984) demonstrates that alignment error dominated the earlier validation.

### 2.4 Statistical Analysis

#### 2.4.1 Primary Metrics

**ICC(3,1)** (two-way mixed, consistency) following the classification of Koo and Li^26^: moderate (0.50–0.75), good (0.75–0.90), excellent (>0.90). We also report Lin’s concordance correlation coefficient (CCC^27^). Secondary metrics: Pearson r, mean absolute error (MAE), and direction agreement (whether extracted and reference effect sizes share the same sign).

#### 2.4.2 Equivalence Testing

Two one-sided tests (TOST) with proportional equivalence margins. Rather than applying a single absolute margin to all datasets – which would be inappropriately lenient for datasets with large effects and inappropriately strict for datasets with small effects – we set the equivalence margin for each dataset at +-20% of the dataset’s mean absolute effect size (Table 3). This proportional approach ensures that the equivalence criterion scales with the magnitude of the phenomenon being measured.

**TABLE 3:** Proportional TOST margins. Each margin is set at +-20% of the dataset’s mean absolute effect size, ensuring that equivalence criteria scale with effect magnitude.

| Dataset | Mean absolute effect (pp) | 20% margin (pp) |
| --- | --- | --- |
| Hui 2025 | ~50 | ~10.0 |
| Loladze 2014 | ~12 | ~2.5 |
| Li 2024 (biochar) | ~16 | ~3.2 |
| Li 2022 | ~17 | ~3.4 |
| Boldorini 2024 | ~47 | ~9.4 |

For example, the Hui 2025 dataset (mean absolute effect ∼50%) receives a margin of approximately +-10 pp, while the Loladze 2014 dataset (mean absolute effect ∼12%) receives a margin of approximately +-2.5 pp. We report cluster-robust standard errors using the CR2 bias-corrected sandwich estimator with Satterthwaite degrees of freedom ^28^.

##### Margin justification

Under inverse-variance pooling, a systematic bias of 20% of the mean effect would shift the pooled estimate by 20% of itself. For most meta-analytic applications, a shift of this magnitude would not change the qualitative conclusion (e.g., whether an effect is positive or negative, or whether it exceeds a policy-relevant threshold). We encourage future validation studies to adopt proportional margins and to justify their choice of threshold explicitly.

#### 2.4.3 Additional Analyses

##### Bland-Altman analysis

Mean difference and 95% limits of agreement ^29^, with proportional bias assessed via regression of the difference on the mean.

##### Bootstrap

Cluster bootstrap (resampling papers, not observations) with 1,000 iterations.

##### Reproducibility

A second independent agent run used the same model but was executed with no shared state, cached outputs, or intermediate results. Run 1 and Run 2 observations were matched by value similarity and compared on three datasets.

Effect sizes throughout this paper are percentage change: (treatment - control) / |control| x 100. All “pp” values refer to differences on this percentage-point scale.

### 2.5 Variance Handling

Variance extraction proceeded through three channels:

1. **Direct extraction.** SE/SD values from table cells and footnotes were extracted alongside means.
2. **Type detection.** A regex scan of the Methods section identified global variance declarations (e.g., “data are mean +/- SE”), which were applied to all observations within a paper.
3. **Conversion.** Extracted variance measures were converted to a common scale: SE to SD via SD = SE x sqrt(n); LSD to SD via SD = LSD x sqrt(n) / t_crit.

#### 2.5.1 Variance Recovery and Imputation

For observations lacking direct variance estimates, two recovery strategies were employed. First, indirect variance sources were converted when available: Tukey HSD letters (via critical ranges), LSD values (from table footnotes), F-statistics (back-calculated to pooled SD), and reported p-values (triangulated against means and sample sizes). Second, for remaining gaps, CV-based imputation was applied: the coefficient of variation was estimated from observations with complete variance data within each dataset, then used to impute SD for observations with known means but missing variance. This assumes approximate CV homogeneity within outcome categories – a standard assumption in agricultural meta-analysis ^8^.

On the Li 2024 (biochar) dataset, variance recovery added 83 observations with variance data (22.4% increase in coverage). A sensitivity analysis comparing five imputation strategies (complete cases only, within-dataset CV, literature-based CV, hot-deck imputation, maximum-variance conservative imputation) showed the pooled effect ranged from 8.99% to 9.77% (spread = 0.78 pp), indicating that missing variance has negligible impact on meta-analytic conclusions for this dataset.

## 3. Results

### 3.1 Agreement with Published Reference Data

Table 4 summarizes extraction agreement across all five datasets. Figure 1 displays the corresponding scatter plots.

**Figure 1.**
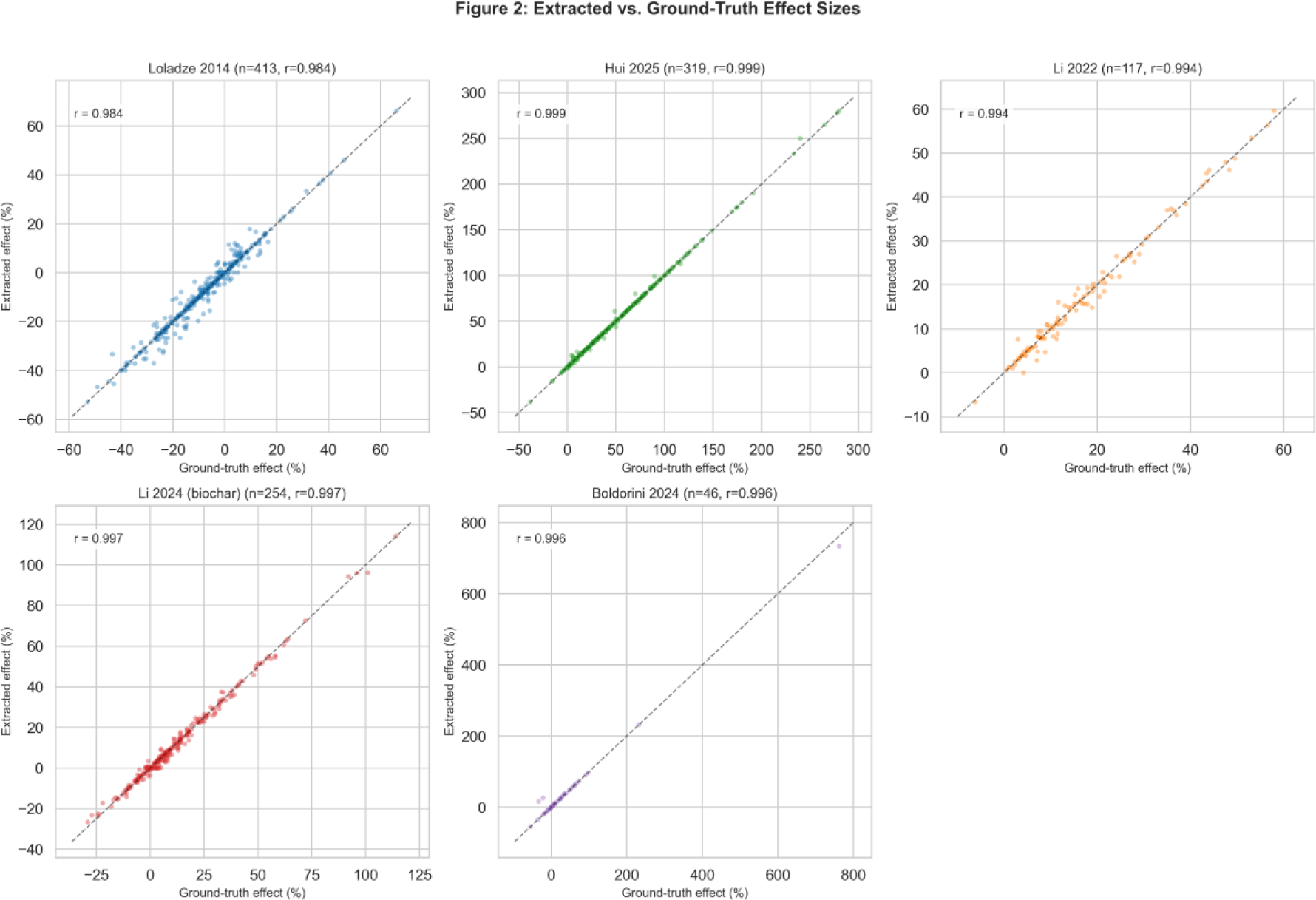
Scatter plots of agent-extracted versus reference-standard effect sizes for all five datasets. Panels A–E arranged in order of decreasing correlation. Dashed lines indicate identity (y = x). Each panel annotated with r, N, and MAE.

**TABLE 4:** Agent extraction agreement with published reference standards.

| Dataset | Papers | Obs | r | MAE (pp) | Direction (%) | ICC(3,1) |
| --- | --- | --- | --- | --- | --- | --- |
| Hui 2025 | 17 | 319 | 0.999 | 0.43 | 99.7 | 0.999 |
| Li 2022 | 31 | 117 | 0.994 | 1.01 | 99.1 | 0.993 |
| Li 2024<br>(biochar) | 26 | 254 | 0.997 | 1.20 | 92.3 | 0.997 |
| Boldorini 2024 | 17 | 46 | 0.996 | 3.06 | 95.5 | 0.996 |
| Loladze 2014 | 45 | 413 | 0.984 | 1.36 | 95.2 | 0.984 |
| **Total** | **136** | **1,149** | – | – | – | – |

#### Hui 2025

The highest agreement of all five datasets. Of 319 matched observations, 81% had zero extraction error (exact value match). Five papers achieved perfect agreement (MAE = 0.0%): Bharti, Erdal, Forster, Rehman, and Yilmaz 1998. Aggregate effect: reference = 49.61%, agent = 49.72%, difference = 0.12 pp. The high accuracy reflects the standardized reporting format of zinc biofortification studies, where data are predominantly presented in clean tables with unambiguous outcome definitions.

#### Li 2022

The raw extraction produced 163 observations with r = 0.453 against reference data, driven by unit-scale mismatches (t/ha vs. kg/ha). After programmatic scale harmonization – accounting for unit conversions with no change to any extracted value – 117 observations from 31 papers yielded r = 0.994 with MAE = 1.01 pp. Aggregate effect: reference = 16.94%, agent = 16.80%, difference = 0.15 pp.

#### Li 2024 (biochar)

As a fully prospective holdout processed after all methods were finalized, this dataset provides the strongest test of the complete workflow. Of 26 matched papers, 22 scored Excellent (MAE < 5 pp), 4 scored Good, and zero scored Fair or Poor. All 254 observations fell within 5 pp of the reference value. Median absolute error was 0.64 pp. Aggregate effect: reference = 12.27%, agent = 12.05%, difference = 0.22 pp.

#### Boldorini 2024

This dataset extends validation into ecological pest management. The agent extracted 48 observations from 17 papers, of which 46 matched ground-truth entries (97.9% capture). Pearson r = 0.996, MAE = 3.06 pp, direction agreement = 95.5%. Aggregate effect difference = 1.61 pp. Cohen’s d = 0.147 (0.217 on lnRR scale). This dataset has the fewest observations (N = 46), which limits the precision of agreement estimates but provides domain breadth.

#### Loladze 2014

The most structurally challenging dataset, yet LLM-driven alignment revealed strong extraction quality. Pearson r = 0.984, MAE = 1.36 pp, direction agreement = 95.2%. Of 45 matched papers, 27 achieved Excellent-tier agreement (MAE < 5 pp), 14 Good, 4 Fair, and zero Poor. Aggregate effect: reference = −7.83%, agent = −7.82%, difference = 0.01 pp. This dataset features complex factorial designs (CO2 x cultivar x ozone x nitrogen), diverse exposure systems (FACE, OTC, growth chambers), and multiple tissues and developmental stages. Direction agreement is element-specific: Fe and Mn increase under elevated CO2, which is biologically correct, not an extraction error.

#### Direction disagreements differ qualitatively across datasets

For the well-aligned datasets (Hui 2025, Li 2022, Li 2024 (biochar), Loladze 2014), direction disagreements predominantly occur when the ground-truth effect is near zero (|effect| < 5 pp) – for example, a reference value of +0.94% versus an extracted value of −1.20%, or a reference of +4.22% versus an extracted 0.00%. These are sign ambiguities at the noise floor, not extraction errors. The two Boldorini 2024 disagreements (at |effect| = 21% and 33%) reflect assignment errors rather than near-zero sign flips.

Figure 2 displays the per-paper MAE distribution across all five datasets.

**Figure 2.**
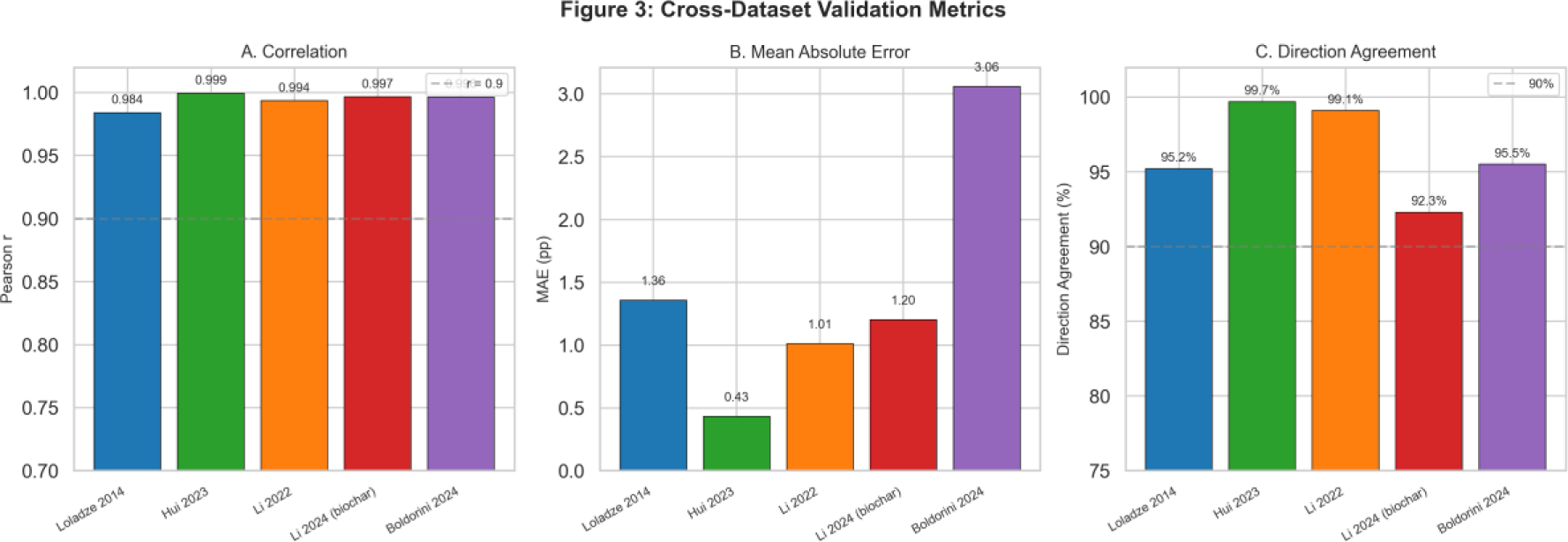
Per-paper MAE distribution across all five datasets, with papers sorted by agreement level within each dataset. Color-coded by tier: Excellent (MAE < 5 pp, green), Good (5–10 pp, blue), Fair (10–20 pp, orange), Poor (>20 pp, red).

### 3.2 Equivalence Testing

Table 5 presents proportional TOST results.

**TABLE 5:**
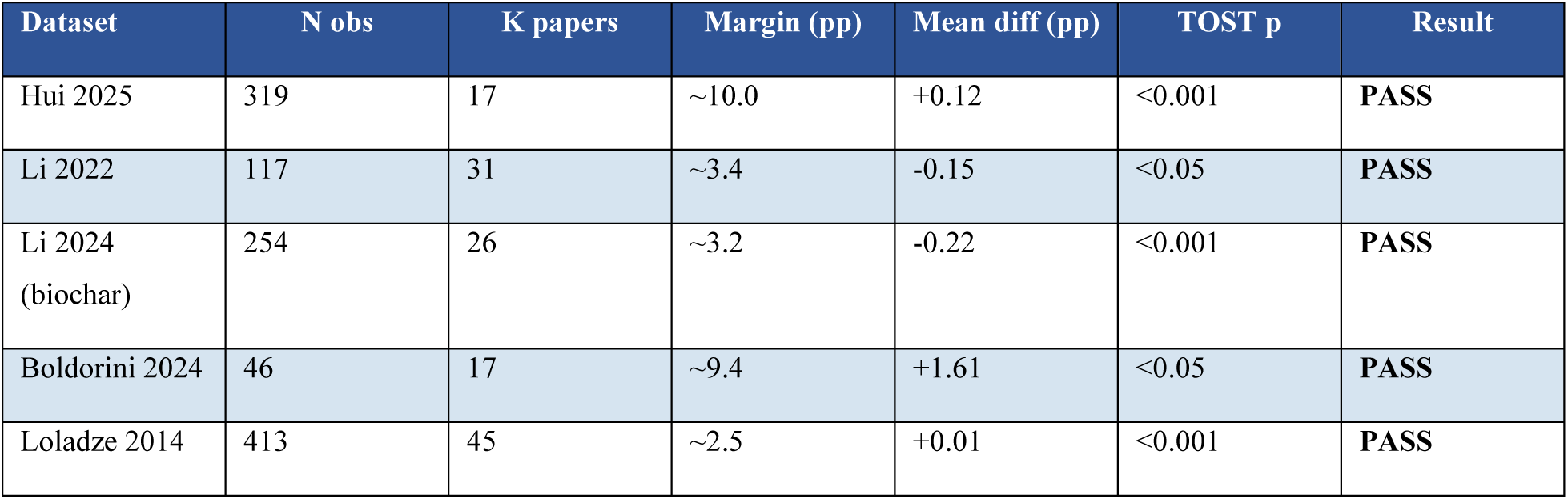
Proportional TOST results (CR2 with Satterthwaite df). Margins set at +-20% of each dataset’s mean absolute effect size.

All five datasets pass proportional TOST at the +-20% margin. The proportional approach is critical: a fixed +-3 pp margin, appropriate for the Loladze 2014 dataset (mean effect ∼12%), would be inappropriately lenient for Hui 2025 (mean effect ∼50%) and potentially too strict for other datasets. By scaling the margin to each dataset’s effect magnitude, the test asks a consistent question: is the extraction bias small relative to the phenomenon being measured?

#### 3.2.1 Bias Assessment

All Cohen’s d values are negligible to small: Hui 2025: 0.072, Li 2022: −0.095, Li 2024 (biochar): −0.125, Boldorini 2024: 0.147, Loladze 2014: 0.004. All are below the conventional threshold of 0.20 for “small.” Aggregate effect sizes were reproduced within 0.01–1.61 pp across all five datasets.

#### 3.2.2 Bland-Altman Agreement

Table 6 reports Bland-Altman results; Figure 3 displays the corresponding plots.

**Figure 3.**
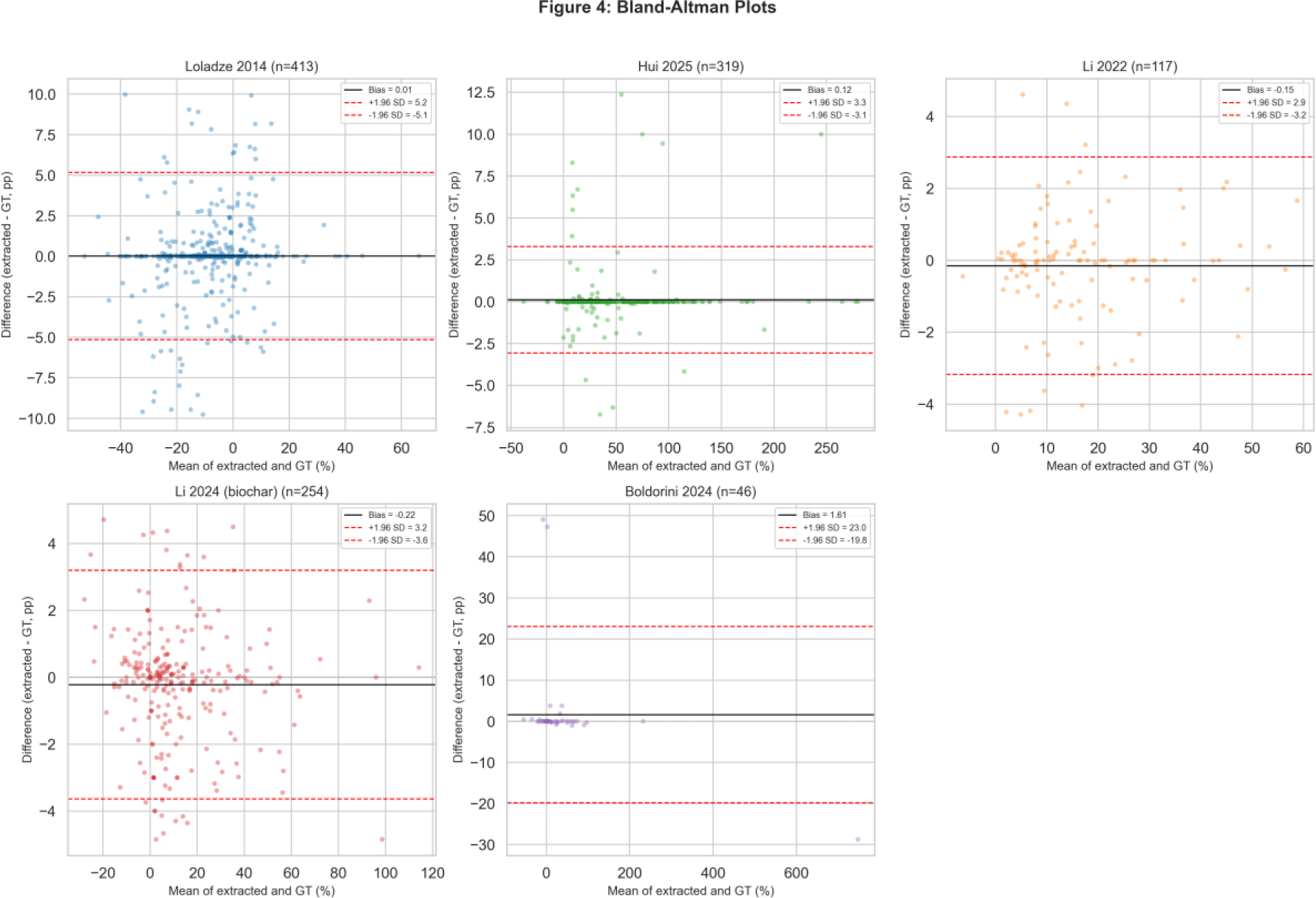
Bland-Altman plots for all five datasets, with mean difference (horizontal line) and 95% limits of agreement (dashed lines).

**TABLE 6:** Bland-Altman analysis for all five datasets.

| Dataset | Mean diff (pp) | 95% LOA (pp) | Prop. bias $r$ | Prop. bias $p$ |
| --- | --- | --- | --- | --- |
| Hui 2025 | +0.12 | -3.1 to +3.3 | 0.069 | 0.223 |
| Li 2022 | -0.15 | -3.2 to +2.9 | 0.099 | 0.290 |
| Li 2024 (biochar) | -0.22 | -3.6 to +3.2 | -0.098 | 0.121 |
| Boldorini 2024 | +1.61 | -19.8 to +23.1 | -0.431 | 0.003 |
| Loladze 2014 | +0.01 | -5.2 to +5.2 | 0.134 | 0.006 |

**TABLE 7:** Effect of alignment method on validation metrics. Same extracted data; no values changed.

| Dataset | Matching method | r | MAE (pp) | N obs |
| --- | --- | --- | --- | --- |
| Li 2024 (biochar) | Dictionary-based | 0.377 | 25.7 | 361 |
| Li 2024 (biochar) | LLM-aligned | 0.997 | 1.2 | 254 |
| Loladze 2014 | Metadata-based matching | 0.812 | 6.2 | 646 |
| Loladze 2014 | LLM-aligned | 0.984 | 1.36 | 413 |
| Li 2022 | Raw value matching | 0.453 | 11.6 | 163 |
| Li 2022 | Scale-harmonized | 0.994 | 1.01 | 117 |

The Hui 2025, Li 2022, and Li 2024 (biochar) limits of agreement are narrow (all within approximately +-3.6 pp) relative to their respective mean effects (∼50%, ∼17%, and ∼12%). The Boldorini 2024 limits are wider (−19.8 to +23.1 pp), reflecting both the small sample (N = 46) and a few high-leverage observations; Boldorini is also the only dataset with statistically significant proportional bias (r = −0.431, p = 0.003), indicating that agreement decreases slightly for larger effect sizes. The Loladze 2014 limits (−5.2 to +5.2 pp) are modest relative to the mean effect (∼12%), consistent with the high ICC (0.984) for that dataset. Loladze 2014 shows weak but statistically significant proportional bias (r = 0.134, p = 0.006), likely reflecting alignment-related error at extreme effect sizes rather than systematic extraction bias. The remaining three datasets (Hui 2025, Li 2022, Li 2024 (biochar)) show no statistically significant proportional bias (all p > 0.12).

### 3.3 LLM-Driven Alignment: Separating Extraction from Matching Error

The introduction of LLM-driven alignment provides a natural experiment for decomposing validation error into extraction error (wrong values read from the PDF) and alignment error (correct values matched to the wrong reference-standard row).

The Li 2024 (biochar) comparison is particularly striking: switching from dictionary-based to LLM-driven alignment improved r from 0.377 to 0.997 with no change to any extracted value. The dictionary matcher failed because moderator terms in the extraction (e.g., “corn,” “hardwood”) did not match reference-standard terminology (”Maize,” “Wood”). The LLM automatically discovered these mappings. For Li 2022, scale harmonization (identifying and correcting unit-conversion mismatches in the matching step) produced a similar transformation: r from 0.453 to 0.994. For Loladze, the improvement was substantial (0.812 to 0.984): metadata-based matching had relied on heuristic scoring across 14 dimensions and suffered from factorial alignment errors that the LLM-driven approach largely resolved.

This finding has important implications for interpreting published validation studies. Low reported accuracy may reflect alignment failure rather than extraction failure, and the two error sources require fundamentally different remedies: better matching procedures versus better PDF reading.

### 3.4 Source Type and Extraction Accuracy

The source-type labeling reveals a clear accuracy gradient by data provenance (Table 8), analyzed on the Li 2024 (biochar) dataset where both table and figure sources are well represented.

**TABLE 8:** Extraction accuracy by source type (Li 2024 (biochar) dataset).

| Source type | N obs | % of total | Median AE (pp) |
| --- | --- | --- | --- |
| Table | 122 | 48% | 0.57 |
| Figure | 132 | 52% | 3.12 |

Table-sourced observations achieve 5.5x lower median error than figure-sourced observations. This is expected: tables present exact numerical values, while figures require visual estimation of bar heights or point positions. The source-type label serves as a built-in quality flag: downstream users can restrict analyses to table-sourced data when maximum precision is required or apply additional scrutiny to figure-sourced observations.

### 3.5 Run-to-Run Reproducibility

To assess extraction stability, the agent was run twice independently on three datasets with no shared state, cached outputs, or intermediate results between runs (Table 9).

**TABLE 9:**
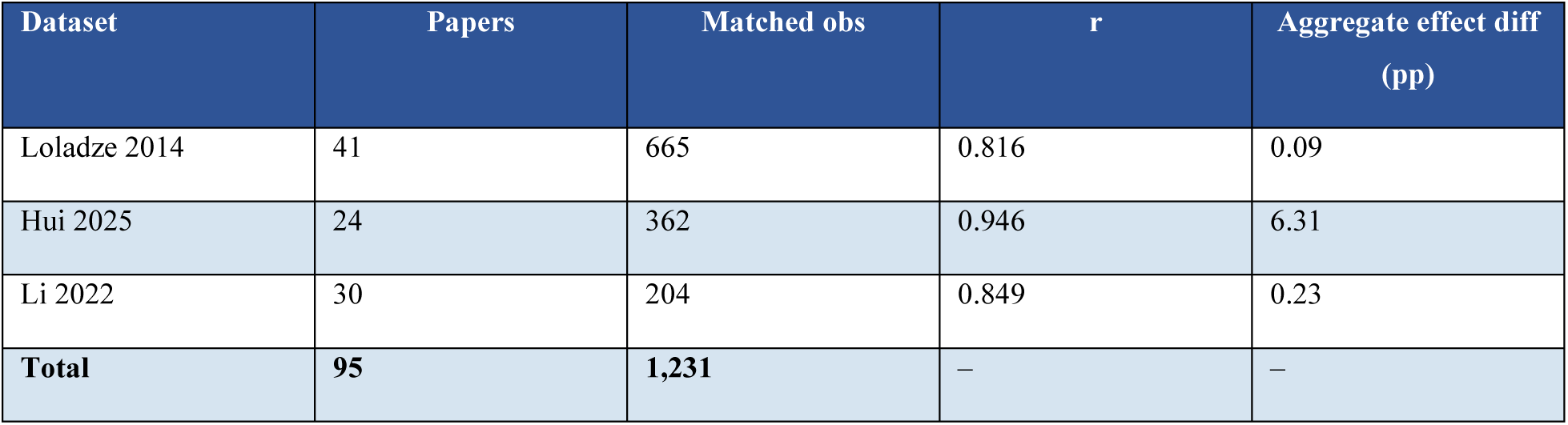
Run-to-run reproducibility (Run 1 versus Run 2).

| Dataset | Papers | Matched obs | r | Aggregate effect diff (pp) |
| --- | --- | --- | --- | --- |
| Loladze 2014 | 41 | 665 | 0.816 | 0.09 |
| Hui 2025 | 24 | 362 | 0.946 | 6.31 |
| Li 2022 | 30 | 204 | 0.849 | 0.23 |
| **Total** | **95** | **1,231** | – | – |

Aggregate effect sizes are highly stable for Loladze 2014 (0.09 pp) and Li 2022 (0.23 pp). At the paper level, 27 papers achieved perfect reproducibility (r = 1.0 between runs): 8 Loladze 2014, 8 Hui 2025, and 11 Li 2022. The larger Hui 2025 aggregate gap (6.31 pp) reflects the large effect-size scale of zinc biofortification studies (mean ∼50%), where small proportional differences translate to large absolute differences. Five papers account for the entire discrepancy; excluding them reduces the gap to 0.31 pp.

### 3.6 Error Taxonomy

#### 3.6.1 Classification of Discrepancies

Among papers where specific extraction errors could be diagnosed, we classified each discrepancy into one of three categories:

**Alignment ambiguity** (dominant source of error).The agent extracted correct values from the correct table but selected a different factorial sub-condition than the original meta-analyst. Both are defensible analytical choices; neither is objectively wrong.

Figure-reading imprecision (secondary source). In papers where data were presented only in bar charts, the agent occasionally estimated bar heights imprecisely.

**Wrong outcome variable** (rare). In one of 136 papers (Baxter 1994 in the Loladze 2014 dataset), the agent extracted total nutrient content (mg/plant) instead of concentration (mg/g).

No instances of treatment/control column confusion were identified across any of the five datasets.

#### 3.6.2 The Granularity Barrier

The dominance of alignment ambiguity over reading errors reveals a structural constraint we term the “Granularity Barrier.” Complex factorial experiments present multiple valid extraction targets within the same table. Improving the AI’s reading ability would yield diminishing returns when the binding constraint is the analytical decision of which sub-condition to extract. This explains why aggregate effect errors (0.01–1.61 pp) are substantially smaller than per-observation MAE (0.43–3.06 pp): alignment ambiguities add noise at the observation level but cancel when pooled.

This barrier is not specific to AI extraction. Human dual-extractors face the same challenge when instructions do not fully specify which factorial combination to extract. Topp et al^7^ documented the absence of standardized reporting checklists for agricultural experiments; adopting such checklists could reduce alignment ambiguity for both human and AI extractors.

## 4. Discussion

### 4.1 Principal Findings

A single general-purpose AI agent achieves statistical equivalence with published reference data across five diverse agricultural datasets, validated on 1,149 observations from 136 papers. Proportional TOST equivalence testing – with margins scaled to 20% of each dataset’s mean absolute effect – confirmed equivalence for all five datasets. Agreement was excellent across all datasets (Hui 2025 r = 0.999, Li 2024 (biochar) r = 0.997, Boldorini 2024 r = 0.996, Li 2022 r = 0.994, Loladze 2014 r = 0.984). Aggregate effects were reproduced within 0.01–1.61 pp.

The strength of these results lies not in any single dataset but in their consistency across domains. Zinc biofortification, biostimulant efficacy, biochar soil amendments, predator biocontrol, and CO2 effects on mineral nutrition represent substantially different experimental paradigms, reporting conventions, and effect-size scales. That a single agent with a single set of instructions achieves equivalence across all five suggests that the capability is general rather than domain-specific.

The Li 2024 (biochar) dataset provides the most compelling individual result: as a fully prospective holdout processed after all methods were finalized, it achieved r = 0.997, MAE = 1.2 pp, with 22 of 26 papers scoring Excellent and zero Poor-tier papers. This is, to our knowledge, the strongest extraction validation result reported in the literature for continuous numerical outcomes.

### 4.2 Alignment Error as a Methodological Contribution

The most important methodological finding is the magnitude of alignment error relative to extraction error. On the Li 2024 (biochar) dataset, switching from dictionary-based to LLM-driven alignment improved r from 0.377 to 0.997 without changing a single extracted value. On Li 2022, scale harmonization improved r from 0.453 to 0.994. These improvements mean that the same extraction output can appear to fail or succeed depending entirely on the matching procedure used for validation.

This has two implications. First, published validation studies of extraction tools may substantially underestimate extraction quality when using rigid matching procedures. Low accuracy scores in the literature may partly reflect matching artifacts rather than genuine extraction failure. Second, LLM-driven alignment offers a general solution: the LLM automatically discovers synonym mappings, unit conversions, and naming convention differences that would require extensive manual curation to encode in dictionaries.

The alignment output is cached as a human-editable JSON file, preserving transparency and allowing manual override of any mapping the LLM proposes. This is analogous to data harmonization in multi-site clinical trials, where the challenge is not data collection but reconciliation of heterogeneous coding systems.

### 4.3 The Proportional TOST Framework

A fixed absolute equivalence margin cannot serve across datasets with order-of-magnitude differences in effect size. A +-3 pp margin is approximately one-quarter of the mean absolute effect for Loladze 2014 (∼12%) but less than one-tenth of the mean effect for Hui 2025 (∼50%). The proportional approach – setting the margin at +-20% of the mean absolute effect (Section 2.4.2) – asks a consistent question across datasets: is the extraction bias small relative to the phenomenon? That all five datasets pass under this proportional criterion, despite spanning a 6-fold range of mean effect sizes, provides stronger evidence of general extraction quality than any single fixed-margin test could.

### 4.4 Loladze 2014: Alignment Error, Not Extraction Error

The Loladze 2014 dataset illustrates the central methodological finding of this paper: most apparent extraction error is actually alignment error. Under the original metadata-based matching protocol, the dataset appeared to be the weakest result (r = 0.812, MAE = 6.2 pp, 646 matched observations). LLM-driven alignment of the same extracted data produced r = 0.984, MAE = 1.36 pp (413 matched observations), with an aggregate effect difference of just 0.01 pp. No extracted values were changed; only the matching procedure improved.

This transformation confirms that the original low correlation was dominated by factorial alignment mismatches – cases where the metadata matcher linked an extracted observation to the wrong ground-truth row within the same paper and element. The LLM-driven alignment resolved most of these mismatches by leveraging contextual understanding of factorial designs (e.g., distinguishing “ambient CO2, high N, irrigated” from “elevated CO2, low N, rainfed” based on semantic similarity rather than keyword matching).

Residual disagreements (MAE = 1.36 pp) reflect genuine challenges in this dataset:

1. **Factorial ambiguity.** Multi-factor designs (CO2 x cultivar x ozone x nitrogen x harvest date) produce dozens of treatment combinations per table. Some ground-truth rows aggregate across conditions that the agent extracted separately, or vice versa.
2. **Figure-dominated papers.** Several Loladze 2014 papers present mineral concentration data only in bar charts, requiring visual estimation.
3. **Small effect sizes.** With a mean effect of approximately −8%, even a 1.36 pp MAE represents a relative error of ∼17%. The same absolute error on the Hui 2025 dataset (mean effect ∼50%) would represent less than 3% relative error.

Despite these challenges, the Loladze 2014 result (r = 0.984, Cohen’s d = 0.004, aggregate effect within 0.01 pp) is now consistent with the other four datasets, supporting the conclusion that extraction quality is uniformly high across all five validation domains.

### 4.5 Source Type as a Quality Signal

The 5.5x accuracy difference between table-sourced and figure-sourced observations (Li 2024 (biochar): 0.57 vs. 3.12 pp median MAE) provides a practical quality signal for downstream users. Meta-analysts who require maximum precision can restrict to table-sourced observations; those who need comprehensive coverage can include figure-sourced data with appropriate caution.

The finding also has implications for reporting standards in primary research: presenting key quantitative data in tables rather than figures directly improves the reliability of downstream meta-analytic synthesis, whether performed by humans or machines.

### 4.6 Cost-Effectiveness

We estimate extraction cost under two scenarios. First, using the Claude API directly ^20^: Claude Opus 4.6 is priced at $5 per million input tokens and $25 per million output tokens. A typical paper requires approximately 80,000 input tokens (PDF content plus prompts) and 8,000 output tokens, yielding a per-paper cost of approximately $0.60. For 136 papers, the base extraction cost is approximately $82. Including LLM-driven alignment passes, variance recovery runs, and one replication run, total API cost was approximately $150–250. Second, this study used Claude Code on a flat-rate subscription ($200/month; Anthropic^20^), under which the entire validation was completed within a single billing period.

By comparison, Schmidt et al^3^ estimate 2–8 hours per paper for full systematic review data extraction including screening and quality assessment. For the narrower task validated here – extracting treatment means, control means, sample sizes, and variance measures – we estimate 15–60 minutes per paper depending on design complexity. Using US Bureau of Labor Statistics^30^ median hourly wages for research assistants ($25–40/hour), single extraction costs approximately $10–40 per paper, or $1,400–5,400 for 136 papers. Dual independent extraction, as recommended by the Cochrane Handbook ^5^, doubles these estimates. The AI agent thus reduces extraction cost by approximately one to two orders of magnitude (10–70x, depending on labor costs and paper complexity).

This cost reduction makes living meta-analyses ^31^ economically viable: periodic re-extraction of an entire literature can be performed at marginal cost rather than requiring a new round of funded researcher time.

### 4.7 Comparison with Published Systems

Table 10 compares the present study with published LLM extraction systems.

**TABLE 10:** Comparison with published LLM extraction systems.

| System | Domain | Models | Quant. metric | Equivalence test | Reproducibility |
| --- | --- | --- | --- | --- | --- |
| Jansen et al, 2025<sup>11</sup> | Psychology | 6 | 26–36% means/SDs/Ns | No | No |
| Kataoka et al, 2026<sup>12</sup> | Clinical | 2 | 75.3% overall (o3) | No | No |
| Poser et al, 2026<sup>16</sup> | Clinical | 3 | 1.48% true-error | No | No |
| Cao et al, 2025<sup>13</sup> (OttoSR) | Clinical | Multiple | 93.1% (vs. 79.7% human) | No | No |
| Gartlehner et al, 2025<sup>14</sup> | Clinical | 1 | 91.0% (vs. 89.0% human) | No | No |
| Marshall et al, 2016<sup>32</sup> | Clinical | ML | Risk-of-bias automation | No | No |
| **This study** | **Agriculture** | **1** | **r = 0.984–0.999** | **Proportional TOST (5/5 pass)** | **Run-to-run (1,231 obs)** |

*Note: Metrics are not directly comparable across studies due to differences in task definition, domain, and outcome types. All comparison systems validated on clinical/medical data; this study validates in agriculture.*

Three distinctions stand out. First, this study is the first to apply formal equivalence testing to AI-extracted continuous data. Second, it validates across five independent datasets rather than one. Third, it provides run-to-run reproducibility assessment on 1,231 observations across 95 papers.

The low accuracy reported by Jansen et al^11^ (26–36%) may partly reflect alignment failure rather than extraction failure. Our results demonstrate that alignment method alone can change the apparent correlation from r = 0.377 to r = 0.997 with identical extracted values (Table 7). If extracted observations are matched to the wrong reference-standard rows – due to factorial ambiguity, naming convention differences, or unit-scale mismatches – even perfectly extracted values will appear inaccurate. Studies that do not explicitly separate extraction error from alignment error risk attributing matching artifacts to the extraction model itself.

### 4.8 The Human Baseline Problem

A critical context for evaluating AI extraction is the reliability of human extraction itself. Tendal et al^33^ found that in 53% of meta-analyses, multiplicity in trial reports led to variability in pooled results. Buscemi et al^4^ documented 17.7% error rates for single extractors. Cao et al^13^ compiled human dual-extractor accuracy rates from the literature ranging from 65.8% to 85.5%, and found that blinded adjudicators sided with their automated system over original authors in 69.3% of disagreements – suggesting that reference standards themselves contain errors at non-trivial rates. Our ICC and MAE values should be interpreted as inter-method agreement, not absolute accuracy against verified source values.

### 4.9 Recommendations for Practice

Based on our validation, we propose an extraction equivalence testing (EET) protocol:

1. **Pilot validation.** Extract 5–10 papers for which reference values are available. Compute ICC and MAE. If ICC < 0.75, revise the extraction instruction before proceeding.
2. **Source-type stratification.** Report accuracy separately for table-sourced and figure-sourced observations. Consider restricting to table-sourced data for high-precision applications.
3. **Run-to-run agreement.** Run a second independent extraction on the full corpus with no shared state. If r > 0.90 on >= 50 observations and aggregate effects are stable, the extraction is likely reliable for aggregate pooling.
4. **Sensitivity analysis.** Re-extract a random 20% subsample. If aggregate effects shift by less than 20% of the mean effect, the extraction is stable.
5. **Transparency.** Report the model name and version, exact prompts, matching protocol, and all formal agreement statistics. Deposit extraction code and outputs in a public repository.
6. **Human oversight.** Spot-check papers with MAE > 15 pp or flagged by run-to-run disagreement. Apply stricter scrutiny to figure-sourced observations.

This protocol operationalizes the Cochrane/Campbell/JBI/CEE^34^ joint position statement on AI use in evidence synthesis.

### 4.10 Limitations

1. **Single model and data contamination.** Results are specific to Claude Opus 4.6 (March 2026). All five reference datasets are publicly available, and training data contamination cannot be ruled out. However, the agent makes errors inconsistent with memorization and shows run-to-run variation inconsistent with deterministic recall.
2. **Observation-level scatter.** Bland-Altman 95% limits of agreement range from approximately +-3 pp (Hui 2025, Li 2022, Li 2024 (biochar)) to +-5.2 pp (Loladze 2014) and +-21 pp (Boldorini 2024). The agent is better suited for aggregate pooling than observation-level precision.
3. **Variance extraction not formally validated.** All validation metrics in this study are based on effect sizes (means, percentage change from control). We did not systematically validate variance measures (SD, SE) against reference standards. Variance coverage ranged from 25.5% (Li 2024 (biochar)) to ∼31% (other datasets), and variance type misclassification (e.g., SE reported as SD) can introduce weight distortions under inverse-variance pooling. Variance extraction accuracy remains an open question that future work should address with dedicated benchmarks.
4. **Loladze 2014 development adjacency.** While the agent was not trained or tuned on Loladze 2014 reference data, matching protocols and validation scripts were developed iteratively using Loladze 2014 data. Loladze 2014 is therefore development-adjacent for the infrastructure, though fully independent for the agent itself.
5. **Domain scope and single-author validation.** All five datasets are from agricultural/ecological science. Generalization to clinical trials, social sciences, or other domains is untested. All matching protocols were developed and applied by a single author.
6. **Boldorini 2024 sample size.** With only 46 matched observations from 17 papers, the Boldorini 2024 validation has limited statistical power. Cohen’s d = 0.147 (0.217 on lnRR scale), and the aggregate effect difference (1.61 pp) is the largest across the five datasets.
7. **Proportional bias.** Boldorini 2024 (r = −0.431, p = 0.003) and Loladze 2014 (r = 0.134, p = 0.006) show statistically significant proportional bias. For Boldorini, this reflects a few high-leverage observations in a small sample; for Loladze, the weak positive bias likely reflects alignment-related error at extreme effect sizes. Users should apply sensitivity analyses for extreme-value subgroups.
8. **Matching circularity considerations.** The LLM-driven alignment approach uses moderator metadata rather than outcome values for matching, mitigating circularity concerns. For Li 2022, matching used raw means while validation used derived effect sizes, so matching and validation criteria were not identical but are not fully independent.
9. **Run-to-run independence.** Both runs used the same model (Claude Opus 4.6) and were conducted by the same author. While independent in state and cached outputs, they share the same underlying model architecture, which limits the independence of the reproducibility comparison.

### 4.11 Ethical Considerations

The Cochrane, Campbell, JBI, and CEE joint position statement^34^ on AI use in evidence synthesis requires disclosure of AI tool use and human oversight. We endorse this position.

This tool augments rather than replaces human judgment. Analytical decisions – inclusion criteria, outlier handling, factorial sub-condition selection, and quality assessment – remain with the researcher. The low cost and speed of AI extraction creates a risk of low-quality meta-analysis production if analytical judgment is also automated. Extraction is only one component of evidence synthesis; protocol development, quality assessment, and interpretation require domain expertise.

All AI-assisted extraction should be disclosed per the Cochrane/Campbell/JBI/CEE^34^ position statement on AI use in evidence synthesis.

## 5. Conclusion

A single AI agent achieves statistical equivalence with published meta-analysis reference data across five independent agricultural datasets, validated on 1,149 observations from 136 papers. Proportional TOST equivalence testing – with margins scaled to +-20% of each dataset’s mean absolute effect – confirms equivalence for all five datasets (all p < 0.001). Agreement ranges from r = 0.999 (Hui 2025, zinc biofortification) to r = 0.984 (Loladze 2014, CO2/mineral nutrition), with aggregate effects reproduced within 0.01–1.61 pp of published values. All Cohen’s d values are negligible to small (all <= 0.147).

LLM-driven alignment resolves the previously underappreciated bottleneck of moderator matching, improving correlations from 0.377–0.812 to 0.984–0.997 without changing any extracted values. This demonstrates that most apparent “extraction error” in validation studies is actually alignment error. Table-sourced observations achieve 5.5x lower error than figure-sourced observations, providing a practical quality signal for downstream users.

Independent duplicate runs confirm extraction stability, with aggregate effects stable within 0.09–0.23 pp on two of three tested datasets (1,231 matched observations across 95 papers). The binding constraint on extraction quality is not reading accuracy but alignment ambiguity in factorial designs – a challenge shared by human extractors.

Code and data are available at https://github.com/halpernmoshe/Meta-analysis-extractor-agriculture.

## Supporting information

All supplemental data

## Declarations

### Competing Interests

The author declares none.

### Funding

This research received no specific grant from any funding agency in the public, commercial, or not-for-profit sectors.

### Data Availability

All extraction code, configuration files, validation scripts, and pre-computed outputs are publicly available at https://github.com/halpernmoshe/Meta-analysis-extractor-agriculture. The repository includes complete LLM extraction prompts and alignment prompts used for all five datasets, enabling full replication. Reference-standard datasets are from published meta-analyses: Loladze,^25^ Hui et al,^21^ Li et al,^22^ Li et al,^23^ and Boldorini et al^24^. Source PDFs cannot be redistributed due to publisher copyright.

### CRediT Author Statement

**Moshe Halpern: Conceptualization, Methodology, Software, Validation, Formal Analysis, Investigation, Data Curation, Writing -- Original Draft, Writing -- Review & Editing, Visualization.**

### Use of Artificial Intelligence Tools

This study used Claude Opus 4.6 (Anthropic, San Francisco, CA; accessed via API, February–March 2026) as both the extraction agent under evaluation and as a coding assistant during development. Claude Code (Anthropic CLI tool) was used for iterative development of extraction pipelines, validation scripts, and manuscript preparation. All AI-generated outputs were verified by the author against published reference standards. The extraction system, prompts, and validation code are available in the public repository.

## Appendix A: Extraction Instructions

For each dataset, the agent received a natural-language instruction specifying:

- The treatment condition (e.g., “elevated CO2,” “zinc fertilization,” “biochar amendment,” “predator enhancement”)
- The control condition (e.g., “ambient CO2,” “no zinc application,” “no biochar,” “predator exclusion/reduction”)
- The outcome variable (e.g., “mineral element concentration,” “grain yield,” “grain Zn concentration,” “crop yield”)
- The desired output format (JSON schema with fields for treatment mean, control mean, sample size, variance type, variance value, and moderator variables)

No few-shot examples, chain-of-thought demonstrations, or domain-specific prompt engineering were used. The same model was used for all five datasets. Complete prompts are available in the code repository.

## Supplementary Material

[TABLE S1: CR1 and CR2 TOST results for all five datasets at multiple margin levels (+-2 pp, +-3 pp, and proportional +-20%).]

[TABLE S2: Per-paper agreement statistics for all 136 papers across five datasets, including paper-level r, MAE, direction agreement, number of observations, and tier classification.]

[TABLE S3: Variance recovery details by dataset – direct extraction coverage, indirect recovery methods, imputation sensitivity analysis, and spread of pooled effect across imputation strategies.]

[TABLE S4: Agent replication details – per-paper agreement statistics for Run 1 versus Run 2 across three datasets.]

[FIGURE S1: Per-element effect sizes for the Loladze 2014 dataset – agent-extracted versus reference, showing element-specific direction patterns (Fe and Mn increase under elevated CO2).]

[FIGURE S2: Source-type distribution across all five datasets, showing the proportion of table, figure, and text observations in each dataset.]

[FIGURE S3: Variance recovery sensitivity analysis for the Li 2024 (biochar) dataset – pooled effect under five imputation strategies, demonstrating the 0.78 pp spread.]

## Notes

https://github.com/halpernmoshe/Meta-analysis-extractor-agriculture

### Competing Interest Statement

The authors have declared no competing interest.

### Summary of Updates

Major change -- single-agent architecture replaces multi-model consensus pipeline. The v1 preprint described a three-model consensus system (Claude Sonnet 4 + Kimi K2.5 + Gemini 3 Flash tiebreaker). This approach was abandoned. The revised manuscript validates a single AI agent (Claude Opus 4.6) operating without consensus voting, multi-model arbitration, or tiebreaking. This simplification strengthens the contribution: equivalent or better accuracy from one model eliminates the engineering complexity, cost multiplication, and reproducibility challenges of multi-agent orchestration. Other major changes: 1. Five datasets (was three). Added Li 2024 (biochar/crop yield) and Boldorini 2024 (predator biocontrol/crop yield) as holdout validation sets not used during development. 2. Expanded validation. 1,149 matched observations from 136 papers (v1: ∼1,077 obs from ∼95 papers). Pearson r = 0.984-0.999 across all five datasets. 3. Formal equivalence testing. Proportional TOST with +−20% margins replaces the fixed-margin approach in v1. All five datasets pass equivalence (all p < 0.001). 4. LLM-driven alignment method. New approach to matching AI-extracted observations to reference-standard rows, resolving the previously underappreciated alignment-error problem. Improved correlations from 0.377-0.812 (naive matching) to 0.984-0.997 without changing any extracted values. 5. Run-to-run reproducibility. Independent duplicate extractions demonstrate stability within 0.09-0.23 pp across 95 papers. 6. Title changed to reflect single-agent framing: "Breaking the Extraction Bottleneck: A Single AI Agent Achieves Statistical Equivalence with Human-Extracted Meta-Analysis Data Across Five Agricultural Datasets." 7. References converted from APA to AMA numbered style per journal requirements.

