## Supplementary material for "Breaking the Extraction Bottleneck: A Single AI Agent Achieves Statistical Equivalence with Human-Extracted Meta-Analysis Data Across Five Agricultural Datasets": All supplemental data

Moshe Halpern

**Table S1. TOST equivalence results for all five datasets at four margin levels (±2 pp, ±3 pp, proportional ±20%, and proportional ±10% of mean absolute effect size). CR2 bias-corrected sandwich estimator with Satterthwaite degrees of freedom.**

| Dataset | Margin_type | Margin_pp | Margin_label | N_obs | Mean_diff_pp | TOST_p | Result |
|---|---|---|---|---|---|---|---|
| Loladze 2014 (mineral/CO2) | Fixed | 2.0 | +/-2pp | 413 | 0.0111 | 0.0 | Equivalent |
| Loladze 2014 (mineral/CO2) | Fixed | 3.0 | +/-3pp | 413 | 0.0111 | 0.0 | Equivalent |
| Loladze 2014 (mineral/CO2) | Proportional (20%) | 2.477 | +/-20% of \|effect\| = +/-2.48pp | 413 | 0.0111 | 0.0 | Equivalent |
| Loladze 2014 (mineral/CO2) | Proportional (10%) | 1.2385 | +/-10% of \|effect\| = +/-1.24pp | 413 | 0.0111 | 0.0 | Equivalent |
| Hui 2025 (Zn/wheat) | Fixed | 2.0 | +/-2pp | 319 | 0.1171 | 0.0 | Equivalent |
| Hui 2025 (Zn/wheat) | Fixed | 3.0 | +/-3pp | 319 | 0.1171 | 0.0 | Equivalent |
| Hui 2025 (Zn/wheat) | Proportional (20%) | 10.0464 | +/-20% of \|effect\| = +/-10.05pp | 319 | 0.1171 | 0.0 | Equivalent |
| Hui 2025 (Zn/wheat) | Proportional (10%) | 5.0232 | +/-10% of \|effect\| = +/-5.02pp | 319 | 0.1171 | 0.0 | Equivalent |
| Li 2022 (biostimulant/yield) | Fixed | 2.0 | +/-2pp | 117 | -0.1472 | 0.0 | Equivalent |
| Li 2022 (biostimulant/yield) | Fixed | 3.0 | +/-3pp | 117 | -0.1472 | 0.0 | Equivalent |
| Li 2022 (biostimulant/yield) | Proportional (20%) | 3.4097 | +/-20% of \|effect\| = +/-3.41pp | 117 | -0.1472 | 0.0 | Equivalent |
| Li 2022 (biostimulant/yield) | Proportional (10%) | 1.7048 | +/-10% of \|effect\| = +/-1.70pp | 117 | -0.1472 | 0.0 | Equivalent |
| Biochar 2024 (biochar/yield) | Fixed | 2.0 | +/-2pp | 254 | -0.2184 | 0.0 | Equivalent |
| Biochar 2024 (biochar/yield) | Fixed | 3.0 | +/-3pp | 254 | -0.2184 | 0.0 | Equivalent |
| Biochar 2024 (biochar/yield) | Proportional (20%) | 3.2181 | +/-20% of \|effect\| = +/-3.22pp | 254 | -0.2184 | 0.0 | Equivalent |
| Biochar 2024 (biochar/yield) | Proportional (10%) | 1.6091 | +/-10% of \|effect\| = +/-1.61pp | 254 | -0.2184 | 0.0 | Equivalent |
| Boldorini 2024 (predator/yield) | Fixed | 2.0 | +/-2pp | 46 | 1.6089 | 0.404745 | Not equivalent |
| Boldorini 2024 (predator/yield) | Fixed | 3.0 | +/-3pp | 46 | 1.6089 | 0.196464 | Not equivalent |
| Boldorini 2024 (predator/yield) | Proportional (20%) | 9.4155 | +/-20% of \|effect\| = +/- | 46 | 1.6089 | 8e-06 | Equivalent |

| | | | | | | | |
|---|---|---|---|---|---|---|---|
| | | | 9.42pp | | | | |
| Boldorini 2024 (predator/yield) | Proportional (10%) | 4.7078 | +/-10% of \|effect\| = +/- 4.71pp | 46 | 1.6089 | 0.030501 | Equivalent |

**Table S2. Per-paper agreement statistics for all papers across five datasets. Tier classification: Excellent (MAE < 5 pp), Good (5–10 pp), Fair (10–20 pp), Poor (> 20 pp).**

| Dataset | Paper_ID | N_obs | MAE_pp | Direction_pct | Tier |
|---|---|---|---|---|---|
| Loladze 2014 | Barnes_1992 | 6 | 0.15 | | Excellent |
| Loladze 2014 | Hogy_2009 | 12 | 1.46 | | Excellent |
| Loladze 2014 | Huluka_1994 | 4 | 6.8 | | Fair |
| Loladze 2014 | Wu_2004 | 4 | 0.0 | | Excellent |
| Loladze 2014 | Keutgen_2001 | 5 | 0.67 | | Excellent |
| Loladze 2014 | Lieffering_2004 | 13 | 4.15 | | Good |
| Loladze 2014 | Pleijel_2009 | 3 | 0.0 | | Excellent |
| Loladze 2014 | Fernando_2012a | 4 | 1.36 | | Excellent |
| Loladze 2014 | 027_Peet_1986 | 5 | 3.19 | | Good |
| Loladze 2014 | 031_Pal_2003 | 4 | 0.0 | | Excellent |
| Loladze 2014 | 032_Kanowski_2001 | 19 | 2.41 | | Good |
| Loladze 2014 | 034_Johnson_2003 | 22 | 0.27 | | Excellent |
| Loladze 2014 | 035_Oksanen_2005 | 9 | 0.0 | | Excellent |
| Loladze 2014 | 036_Schenk_1997 | 27 | 0.07 | | Excellent |
| Loladze 2014 | 037_Haase_2008 | 1 | 3.07 | | Good |
| Loladze 2014 | Al-Rawahy_2013 | 7 | 2.22 | | Good |
| Loladze 2014 | Azam_2013 | 36 | 0.44 | | Excellent |
| Loladze 2014 | Baslam_2012 | 10 | 2.35 | | Good |
| Loladze 2014 | Baxter_1994 | 1 | 9.59 | | Fair |
| Loladze 2014 | Blank_2011 | 1 | 2.05 | | Good |
| Loladze 2014 | Campbell_2002 | 1 | 9.97 | | Fair |
| Loladze 2014 | Fangmeier_2002 | 24 | 1.59 | | Excellent |
| Loladze 2014 | Fernando_2012 | 8 | 0.32 | | Excellent |
| Loladze 2014 | Finzi_2001 | 10 | 0.23 | | Excellent |
| Loladze 2014 | Guo_2011 | 5 | 1.56 | | Excellent |
| Loladze 2014 | Heagle_1993 | 7 | 3.03 | | Good |
| Loladze 2014 | Housman_2012 | 7 | 0.64 | | Excellent |
| Loladze 2014 | Khan_2013 | 18 | 0.53 | | Excellent |
| Loladze 2014 | Luomala_2005 | 7 | 3.72 | | Good |
| Loladze 2014 | Mjwara_1996 | 3 | 3.21 | | Good |
| Loladze 2014 | Natali_2009 | 16 | 1.65 | | Excellent |
| Loladze 2014 | Newbery_1995 | 2 | 3.0 | | Good |
| Loladze 2014 | Niinemets_1999 | 9 | 0.53 | | Excellent |
| Loladze 2014 | Niu_2013 | 2 | 1.25 | | Excellent |
| Loladze 2014 | ONeill_1987 | 12 | 0.02 | | Excellent |
| Loladze 2014 | Overdieck_1993 | 22 | 0.79 | | Excellent |

| | | | | | |
|---|---|---|---|---|---|
| Loladze 2014 | Pfirrmann_1996 | 6 | 3.9 | | Good |
| Loladze 2014 | Polley_2011 | 5 | 3.4 | | Good |
| Loladze 2014 | Porter_1984 | 5 | 0.0 | | Excellent |
| Loladze 2014 | Rodenkirchen_2009 | 13 | 4.01 | | Good |
| Loladze 2014 | Seneweera_1997 | 10 | 1.18 | | Excellent |
| Loladze 2014 | Singh_2013 | 10 | 0.17 | | Excellent |
| Loladze 2014 | Wilsey_1994 | 13 | 0.03 | | Excellent |
| Loladze 2014 | Woodin_1992 | 3 | 5.91 | | Fair |
| Loladze 2014 | Ziska_1997 | 2 | 0.02 | | Excellent |
| Hui 2025 | (aggregate - 319 obs) | 319 | 0.43 | | Excellent |
| Li 2022 | 009_Ali_2019_Biostimulatory activities of Ascophyllum nodosum e | 1 | 0.54 | 100.0 | Excellent |
| Li 2022 | 027_Chen_2021_Effects of Seaweed Extracts on the Growt | 3 | 0.18 | 100.0 | Excellent |
| Li 2022 | 029_Ciepiela_2019_The effect of biostimulants derived from | 3 | 1.35 | 100.0 | Excellent |
| Li 2022 | 058_Fichhof_2018_Management of Biostimulant and Silicon i | 1 | 4.29 | 100.0 | Excellent |
| Li 2022 | 067_Grabowska_2012_The Effect of Cultivar and Biostimulant | 2 | 0.48 | 100.0 | Excellent |
| Li 2022 | 086_Knapowski_2019_Crop stimulants as a factor determining | 7 | 1.14 | 100.0 | Excellent |
| Li 2022 | 088_Kocira_2019_Effect of amino acid biostimulant on the | 8 | 1.18 | 100.0 | Excellent |
| Li 2022 | 090_Kocira_2020_Biochemical and economical effect of app | 4 | 0.65 | 100.0 | Excellent |
| Li 2022 | 091_Kocira_2018_Modeling biometric traits, yield and nut | 3 | 0.99 | 100.0 | Excellent |
| Li 2022 | 094_Kowalska_2021_Effect of Different Forms of Silicon on | 1 | 1.13 | 100.0 | Excellent |
| Li 2022 | 095_Kuisma_1989_The effect of foliar application of seaw | 2 | 1.43 | 100.0 | Excellent |
| Li 2022 | 1-s2.0-S0304423819306703-main | 6 | 0.01 | 100.0 | Excellent |
| Li 2022 | 1-s2.0-S0304423820302417-main | 10 | 1.37 | 90.0 | Excellent |
| Li 2022 | 1-s2.0-S1878818119307637-main | 12 | 0.0 | 100.0 | Excellent |
| Li 2022 | 1-s2.0-S1878818119309879-main | 1 | 3.22 | 100.0 | Excellent |
| Li 2022 | 105_Mattner_2018_Increased growth response of strawberry | 3 | 1.41 | 100.0 | Excellent |
| Li 2022 | 106_Matysiak_2018_Herbicides with natural and synthetic bi | 1 | 0.26 | 100.0 | Excellent |
| Li 2022 | 110_Michalak_2016_Evaluation of supercritical extracts of | 3 | 1.32 | 100.0 | Excellent |
| Li 2022 | 116_Nurdiawati_2019_Liquid feather protein hydrolysate as a | 1 | 0.45 | 100.0 | Excellent |
| Li 2022 | 120_Pohl_2019_The Eggplant Yield and Fruit Composition | 3 | 1.4 | 100.0 | Excellent |
| Li 2022 | 127_Radkowski_2018_Influence of foliar fertilization with a | 8 | 1.0 | 100.0 | Excellent |
| Li 2022 | 131_Rahman_2018_Chitosan biopolymer promotes yield and s | 6 | 1.02 | 100.0 | Excellent |
| Li 2022 | 1542-1558-15(3)2018 BR-18-165 | 2 | 2.15 | 100.0 | Excellent |
| Li 2022 | 158_Sulakhudin_2019_Application of Coastal Sediments and Fol | 4 | 0.37 | 100.0 | Excellent |
| Li 2022 | 175_Wilczewski_2018_Response of sugar beet to humic substanc | 2 | 3.3 | 100.0 | Excellent |
| Li 2022 | 604-615-14(3)2017BR-1503 | 1 | 1.47 | 100.0 | Excellent |

| | | | | | |
|---|---|---|---|---|---|
| Li 2022 | agriculture-10-00618-v2 | 4 | 2.34 | 100.0 | Excellent |
| Li 2022 | ali | 2 | 0.76 | 100.0 | Excellent |
| Li 2022 | article1400838000_Azarpour et al | 4 | 0.95 | 100.0 | Excellent |
| Li 2022 | plants-09-01633 | 8 | 0.83 | 100.0 | Excellent |
| Li 2022 | sustainability-11-02171 | 1 | 1.66 | 100.0 | Excellent |
| Biochar 2024 | 001_Adekiya_2019 | 4 | 0.08 | | Excellent |
| Biochar 2024 | 007_Gathorne-Hardy_2009 | 1 | 2.0 | | Excellent |
| Biochar 2024 | 016_Li_B_2016 | 6 | 1.56 | | Excellent |
| Biochar 2024 | 021_Nobile_2022 | 12 | 1.22 | | Excellent |
| Biochar 2024 | 041_Guerena_2013 | 12 | 0.89 | | Excellent |
| Biochar 2024 | 063_Asai_2009 | 15 | 1.01 | | Excellent |
| Biochar 2024 | 077_Zhang_J_2019 | 11 | 1.8 | | Excellent |
| Biochar 2024 | 078_Wang_2012 | 14 | 1.35 | | Excellent |
| Biochar 2024 | 081_Deenik_2010 | 8 | 1.63 | | Excellent |
| Biochar 2024 | 082_Jose_2013 | 7 | 1.68 | | Excellent |
| Biochar 2024 | 101_Liang_Feng_2014 | 9 | 0.32 | | Excellent |
| Biochar 2024 | 116_Farrell_2014 | 12 | 1.98 | | Excellent |
| Biochar 2024 | 130_Azeem_2019 | 8 | 0.48 | | Excellent |
| Biochar 2024 | 133_Pandit_2018 | 14 | 2.02 | | Good |
| Biochar 2024 | 145_Omara_2020 | 6 | 2.63 | | Good |
| Biochar 2024 | 153_Wei_2022 | 4 | 2.11 | | Good |
| Biochar 2024 | 166_Haefele_2011 | 20 | 0.98 | | Excellent |
| Biochar 2024 | 184_Yeboah_2018 | 7 | 1.46 | | Excellent |
| Biochar 2024 | 193_Islami_2011 | 9 | 0.67 | | Excellent |
| Biochar 2024 | 207_Liu_2019 | 22 | 1.11 | | Excellent |
| Biochar 2024 | 219_Xie_2021 | 12 | 0.18 | | Excellent |
| Biochar 2024 | 223_Dong_2019 | 2 | 1.67 | | Excellent |
| Biochar 2024 | 227_Niu_2017 | 9 | 1.37 | | Excellent |
| Biochar 2024 | 229_Shi_2022 | 17 | 0.26 | | Excellent |
| Biochar 2024 | 231_Zhang_2021 | 7 | 1.48 | | Excellent |
| Biochar 2024 | 242_Liu_2014 | 6 | 2.33 | | Good |
| Boldorini 2024 | Ali | 1 | 1.01 | 100.0 | Excellent |
| Boldorini 2024 | Bisseleua | 1 | 0.03 | 100.0 | Excellent |
| Boldorini 2024 | Borkhataria | 1 | 0.0 | 100.0 | Excellent |
| Boldorini 2024 | Classen | 1 | 0.16 | 100.0 | Excellent |
| Boldorini 2024 | Garfinkel | 3 | 0.06 | 100.0 | Excellent |
| Boldorini 2024 | Gras | 2 | 0.07 | 100.0 | Excellent |
| Boldorini 2024 | Hooks_et_al | 4 | 0.0 | 100.0 | Excellent |
| Boldorini 2024 | Ismoilov | 1 | 0.0 | 100.0 | Excellent |
| Boldorini 2024 | Lang | 9 | 0.0 | 100.0 | Excellent |
| Boldorini 2024 | Libran-Embid | 1 | 0.0 | 100.0 | Excellent |
| Boldorini 2024 | Maas | 2 | 48.16 | 0.0 | Poor |
| Boldorini 2024 | Martin | 1 | 0.04 | 100.0 | Excellent |
| Boldorini 2024 | Mols | 1 | 0.0 | 100.0 | Excellent |
| Boldorini 2024 | Saunders | 1 | 0.0 | 100.0 | Excellent |
| Boldorini 2024 | Snyder_Wise | 12 | 2.8 | 100.0 | Excellent |
| Boldorini 2024 | Suenaga_Hamamura | 4 | 1.35 | 100.0 | Excellent |
| Boldorini 2024 | Vichitbandha_Wise | 1 | 3.77 | 100.0 | Excellent |

**Table S3. Variance recovery details by dataset, including direct extraction coverage, indirect recovery, and imputation sensitivity.**

| Dataset | N_matched | Direct_variance_pct | Indirect_recovery_count | Indirect_recovery_pct | Imputation_spread_pp | Table_obs | Figure_obs | Text_obs | Table_MAE | Figure_MAE | Notes |
|---|---|---|---|---|---|---|---|---|---|---|---|
| Biochar 2024 | 254 | 25.5 | 83 | 22.4 | 0.78 | 67 | 133 | 10 | 0.61 | 1.41 | Table data 5.5x more precise than figure data |
| Loladze 2014 | 413 | N/A | N/A | N/A | N/A | | | | | | GT uses percentage change; variance not validated separately |
| Hui 2025 | 319 | N/A | N/A | N/A | N/A | | | | | | Validated on effect sizes; variance not separately assessed |
| Li 2022 | 117 | N/A | N/A | N/A | N/A | | | | | | Effect-size-only validation |
| Boldorini 2024 | 46 | N/A | N/A | N/A | N/A | | | | | | lnRR-based validation |

**Table S4. Agent replication stability. Aggregate effect difference between independent duplicate runs.**

| Datase | N_matched_ob | N_paper | Aggregate_effect_Run | Aggregate_effect_Run | Effect_diff_p | Notes |
|---|---|---|---|---|---|---|

| t | s | s | 1 | 2 | p | |
|---|---|---|---|---|---|---|
| Loladze 2014 | 665 | 41 | -4.95% | -5.04% | 0.09 | Run1 vs Run2 agent extraction |
| Hui 2025 | 362 | 24 | | | 6.31 | Large effect-size scale amplifies small proportional differences |
| Li 2022 | 204 | 30 | 10.16% | 10.39% | 0.23 | Aggregate effect stable across runs |

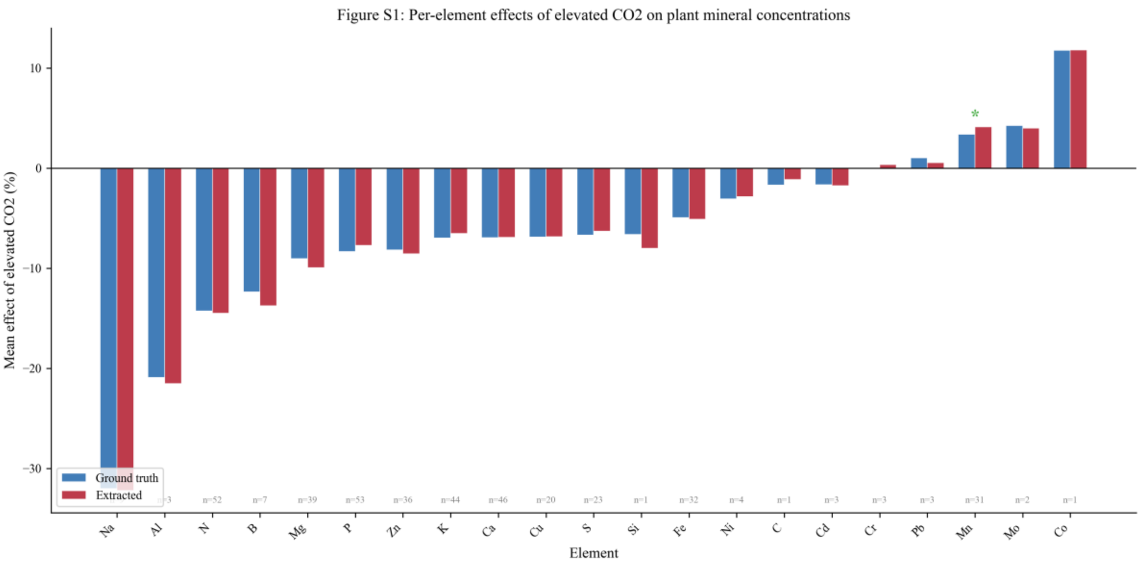


*Figure S1. Per-element effect sizes for the Loladze 2014 dataset. Agent-extracted (orange) versus reference (blue) mean percentage change under elevated $CO_2$. Elements marked with * (Fe, Mn) increase under elevated $CO_2$, which is biologically correct. Error reflects alignment and extraction combined.*

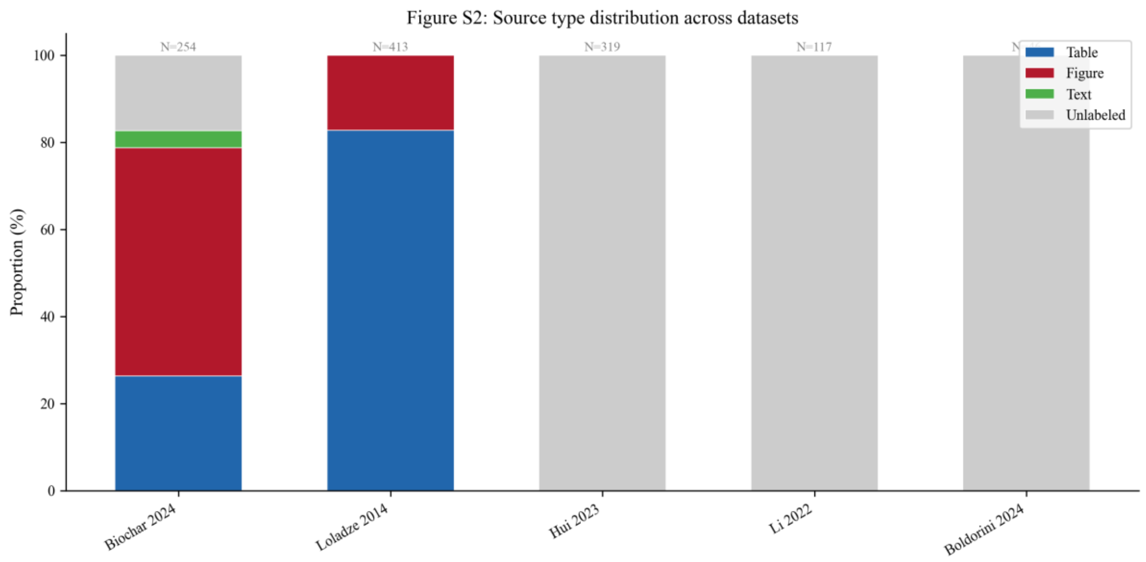


*Figure S2. Source-type distribution across datasets. The Li 2024 (biochar) dataset has detailed source labeling; other datasets show available classification.*

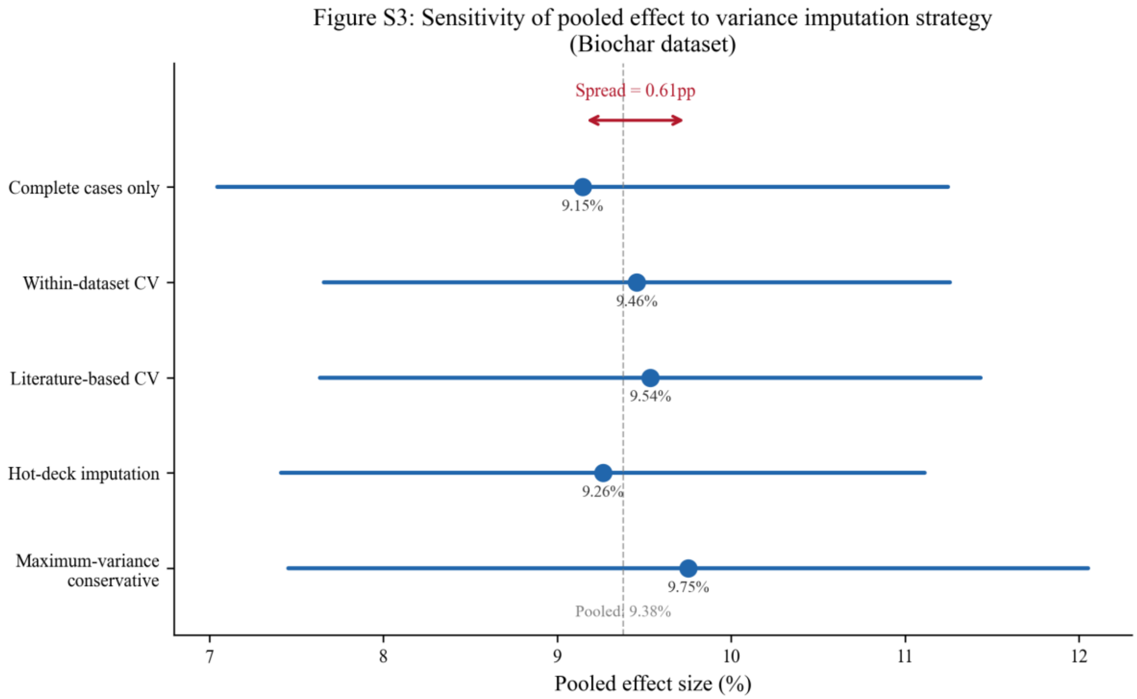


*Figure S3. Variance recovery sensitivity analysis for the Li 2024 (biochar) dataset. Pooled effect size under five imputation strategies, demonstrating robustness (spread < 1 pp).*
